# Mental Effort and Counterfactuals Modulate Language Understanding: ERP Evidence in Older Adults

**DOI:** 10.1101/2024.09.30.612291

**Authors:** José Luis Salas-Herrera, Mabel Urrutia Martínez, Nicolás Andrés Hinrichs

## Abstract

The relationship between language and physical effort in older adults is a field that is scarcely explored in the literature associated with embodiment. An electrophysiological experiment was conducted to explore the modulation of two linguistic contexts: factual and counter-factual, in relation to physical and mental effort using electrophysiological components. 27 older adults (M = 70.34 years, SD = 4.82, 15 women and 12 men) read sentences on a computer screen and responded to an activation test. The results indicate that the linguistic, factual, and counterfactual contexts, as well as the embodiment parameter of mental effort modulate the understanding of language and participate with variable preponderance in different time windows. Furthermore, counterfactuality seems to facilitate the processing of high mental effort, and both factual and counterfactual language elicit the N400 component. These findings contribute to the growing body of research on embodied cognition by providing novel insights into the nuances of cognitive demands involved in language processing in aging population, paving the way for developing targeted interventions aimed at improving communication and cognitive well-being in older adults.

## 1. Introduction

Embodied cognition suggests that language comprehension isn’t just an abstract process that occurs in the brain, but also involves sensorimotor simulations. For example, while reading or hearing the sentence “John kicked the ball”, these words aren’t processed only at a linguistic level, but brain areas related to physical movement are triggered and engaged, too (as if visualizing or imagining the action of to kick a ball). Moreover, theories of embodiment posit that meaning is intimately embedded in the brain’s perceptual, motor, and emotional systems [1, 2, 3, 4]. From an embodied perspective, the body and the specific cerebral cortices related to sensation, motor skills, and emotions have varying degrees of participation in a dimension that ranges from the total embodiment of semantic processing and representation to full disembodiment. The idea that sensory and motor experiences form the basis of conceptual knowledge has a long history in philosophy, psychology, and neuroscience [5, 6, 7, 8]. In recent years, this proposal has gained new steam under the rubric of ‘embodied’ or ‘situated’ cognition, supported by numerous theoretical, neuroimaging, and behavioral studies [9, 10, 11, 12]. Recent years have witnessed the rise of embodied cognition approaches that delve deeper into the embodied essence of language processing [13]. These approaches illuminate the intricate interplay between bodily experiences, sensorimotor processes, and linguistic representations, elucidating how embodiment influences language comprehension and production in real-time scenarios [14]. Moreover, the amalgamation of predictive processing frameworks with embodiment theories has spurred investigations into how the brain formulates and revises predictions through embodied simulations, enriching our comprehension of anticipation’s role in language processing [15]. This integration has significantly advanced our understanding of how the brain processes language dynamically and refines predictions based on embodied simulations [16] with different sensorimotor parameters. Specifically, in the parameter of physical effort, psycholinguistic research had investigated the relationship between language and physical effort [17] on topics such as the relationship between physical effort and counterfactual sentences [18], or the ones between balance and posture, and their spatial and temporal representations [19]. Furthermore, many metaphors and idiomatic expressions have also been shown to reflect physical and sensory experiences, like “grasping a concept” or “shouldering a burden” [1, 2, 20, 3]. The activation of motor representations would be directly related to action verb processing [21]. Since the motor plans caused by the verbs must be integrated with the properties of the associated objects, we suppose that aging could affect the estimation of the direct object, and thus, sentence reading when this implies variable levels of effort. This is because the abilities to perform physical effort in older people are impaired compared to young people [22]. In fact, several studies have shown that older people have difficulties using imaging and motor simulation [23]. Simulations utilize embodiment or neural parameters, such as direction for reaching and force/physical effort for grasping an object, imposing a hierarchical structure on the brain [24]. Just et al. [25] found that low-effort imagery activates the left temporal cortex and primary visual cortex, while high-effort imagery requires generating a multidimensional internal structure of the event. Urrutia et al. [18] identified brain areas sensitive to effort in action sentences, overlapping with regions activated during physical effort tasks. These areas include the left inferior parietal lobe, supramarginal and postcentral areas, associated with action planning. Prefrontal areas were exclusively activated for counterfactual sentences. Action language involves events performed by animate beings and entails physical effort. Research predominantly focuses on physical content involving hand and arm movements [17], but studies have also examined mouth and leg movements [26, 3, 12].

### 1.1. Mental Effort

Mental effort, or mental workload, has been defined as the amount of cognitive resources expended to perform a task, with more complex tasks requiring greater mental effort [27]. Using fMRI, ERP, and pupillometry measures, it has been shown that increased cognitive load or mental effort is often associated with increased activation in specific brain regions such as the prefrontal cortex (PFC) [28]. In assessments of mental effort and workload using fMRI, activation of neural networks such as the executive control network (ECN) and deactivation of the default mode network (DMN) were found under conditions of high cognitive load [29]. Specific EEG and ERP studies have found that ERP amplitudes decrease with increasing mental workload, indicating greater cognitive effort [30]. Variations in ERP components related to task difficulty have been observed; results show that early and late ERP components vary with difficulty, suggesting an increase in cognitive load with more difficult tasks [31]. Similarly, Shaw and colleagues [32] demonstrated that cognitive workload was inversely proportional to attentional reserve. Significant changes in theta and alpha band power have been observed, especially in the prefrontal, frontal, central, temporal, parietal, and occipital regions of the brain. These changes in brain activity have been correlated with different levels of mental workload during working memory tasks, such as the n-back task; specifically, an increase in theta band power and a decrease in alpha band power were found to be associated with increased engagement in more complex tasks [33].

### 1.2. Greater Mental Effort in Counterfactual Stories

Cognitively, understanding counterfactuals involves constructing and maintaining multiple mental representations: the actual state of the world and the hypothetical state posited by the sentence [34, 26]. This requires additional processes of integration and conflict management between representations compared to factual constructions. This cognitive complexity has been investigated through various methodological approaches, which have shed light on the mechanisms underlying counterfactual processing. Behavioral studies have shown that counterfactuals involve longer reading times and poorer subsequent recall than their factual counterparts, suggesting greater cognitive demand [26, 35]. In particular, De Vega et al. [26] found that reading counterfactual sentences embedded in stories resulted in longer reading and reinsertion times of prior information compared to factual ones, reflecting more demanding integration processes. These psycholinguistic findings establish different processing profiles for factual and counterfactual language.

Complementing the above, neuroimaging and electrophysiology research has provided valuable data on counterfactual processing. Brain regions associated with cognitive control, such as the dorsolateral prefrontal cortex and anterior cingulate, have shown increased activation during counterfactual processing [36, 18], supporting the notion that they involve greater mental effort. This extra effort is manifested not only in brain activation, but also in physiological indices of cognitive load. Furthermore, event-related potential (ERP) studies have revealed distinctive patterns in counterfactual processing. Brain potentials such as the N400 have been shown to be sensitive to violations of expectations based on world knowledge in counterfactual contexts [35, 37, 18]. These electrophysiological findings provide further evidence for the specific neural mechanisms recruited during counterfactual information processing. Taken together, research approaches converge in supporting the notion that counterfactual processing is cognitively more demanding and recruits distinctive mechanisms compared to factual language. Our hypothesis is that processing counterfactual sentences requires greater mental effort than factual sentences because counterfactuals involve maintaining multiple mental representations. This increased cognitive load demands additional processes like integration, conflict management, and inhibition of incompatible information, leading to more pronounced electrophysiological responses (e.g., N400 wave). Additionally, we expect that the higher cognitive demands of processing abstract language in counterfactual contexts will result in larger N400 amplitudes and potentially longer latencies, particularly in older adults, compared to the processing of concrete language in indicative contexts. This increased N400 effect would reflect the greater difficulty in semantic integration and the recruitment of additional cognitive resources necessary for counterfactual comprehension in aging populations.

## 2. Materials and Methods

### 2.1. Participants

The sample consisted of 27 older adults from Concepción, Chile (M = 70.34 years, SD = 4.82; 15 women and 12 men). According to the Edinburgh Handedness Inventory [38], all participants were right-handed with a laterality coefficient above 60. One participant was excluded due to excessive artifacts in EEG recordings. All participants passed psychological tests and showed no central nervous system issues, substance abuse, learning or memory problems, mental health issues, serious medical illnesses, or medications affecting the central nervous system. All had normal or corrected vision and were native Spanish speakers. Before the experiment, each participant was evaluated with cognitive tests, screening, and psychopathological scales to exclude participants with conditions such as depression or dementia. The Mini-Mental State Examination (MMSE) [39] was used with a sensitivity of 93.6% and specificity of 46.1% [40]. The Yesavage Depression Scale [41] was also applied, with internal consistency and construct reliability ensuring the reliability of the measures.

### 2.2. Design

The experimental conditions were analyzed using a factorial design with two types of contexts (factual and counterfactual) and two levels of mental effort (low and high).

### 2.3. Materials and Procedures

Linguistic experts (2) and the researcher (psychologist) created over 400 sentences with varying content of mental effort. A normative study involved 25 older people, who rated each sentence for the level of perceived mental effort using a Likert scale. When selecting and evaluating the sentences, relevant embodied aspects such as the imaginability of the actions and congruence with the participants’ sensorimotor experiences were considered to ensure that the selected sentences accurately reflected the mental effort involved.

Based on the results of the normative study, the sentences were categorized into two groups: high and low mental effort. This step involved a statistical analysis of the scores assigned to each sentence, allowing for the identification of those with a clear distinction in the perception of effort. Sentences with intermediate or ambiguous scores between high and low mental effort were eliminated or not considered. An internal consistency analysis was performed to ensure that sentences categorized as high or low mental effort were consistently rated across different participants, thus increasing the validity of the selected stimulus set. The selected sentences were reviewed again by the research team and the linguistics experts for final adjustments. This included a detailed consideration of pragmatic and corporeal aspects, such as the relevance and applicability of the described actions in the everyday contexts of the participants.

A total of 160 sentences corresponding to four types of semantics were elaborated: 40 high mental effort factual sentences (HFS), 40 low mental effort factual sentences (LFS), 40 high mental effort counterfactual sentences (HCS), and 40 low mental effort counterfactual sentences (LCS). These sentences are exemplified in Table 1. Participants began the task by viewing dashes and spaces on the screen, representing the letters of words and sentence segments (see Figure 1 below). Pressing a key revealed the next segment and hid the previous one. After each sentence, they determined whether a test word was present in the sentence by pressing P (marked with a red sticker) for “yes” or Q (marked with a green sticker) for “no.”

**Table 1.** Examples of stimuli in the reading task.

| Item | Sentence | Spanish translation | Condition |
| --- | --- | --- | --- |
| LFS | NOW YOU ARE COUNTING THE TRUCKS ON THE ROAD | AHORA ESTÁS CONTANDO LOS CAMIONES EN LA AVENIDA | Low Mental Effort Factual Sentence |
| HFS | NOW YOU ARE PLANNING THE STRATEGY FOR THE MATCH | AHORA ESTÁS PLANEANDO LA ESTRATEGIA PARA EL PARTIDO | High Mental Effort Factual Sentence |
| LCS | YOU SHOULD HAVE COUNTED THE TRUCKS ON THE AVENUE | DEBERÍAS HABER CONTADO LOS CAMIONES EN LA AVENIDA | Low Mental Effort Counterfactual Sentence |
| HCS | YOU SHOULD HAVE PLANNED THE STRATEGY FOR THE MATCH | DEBERÍAS HABER PLANEADO LA ESTRATEGIA PARA EL PARTIDO | High Mental Effort Counterfactual Sentence |

**Figure 1.**
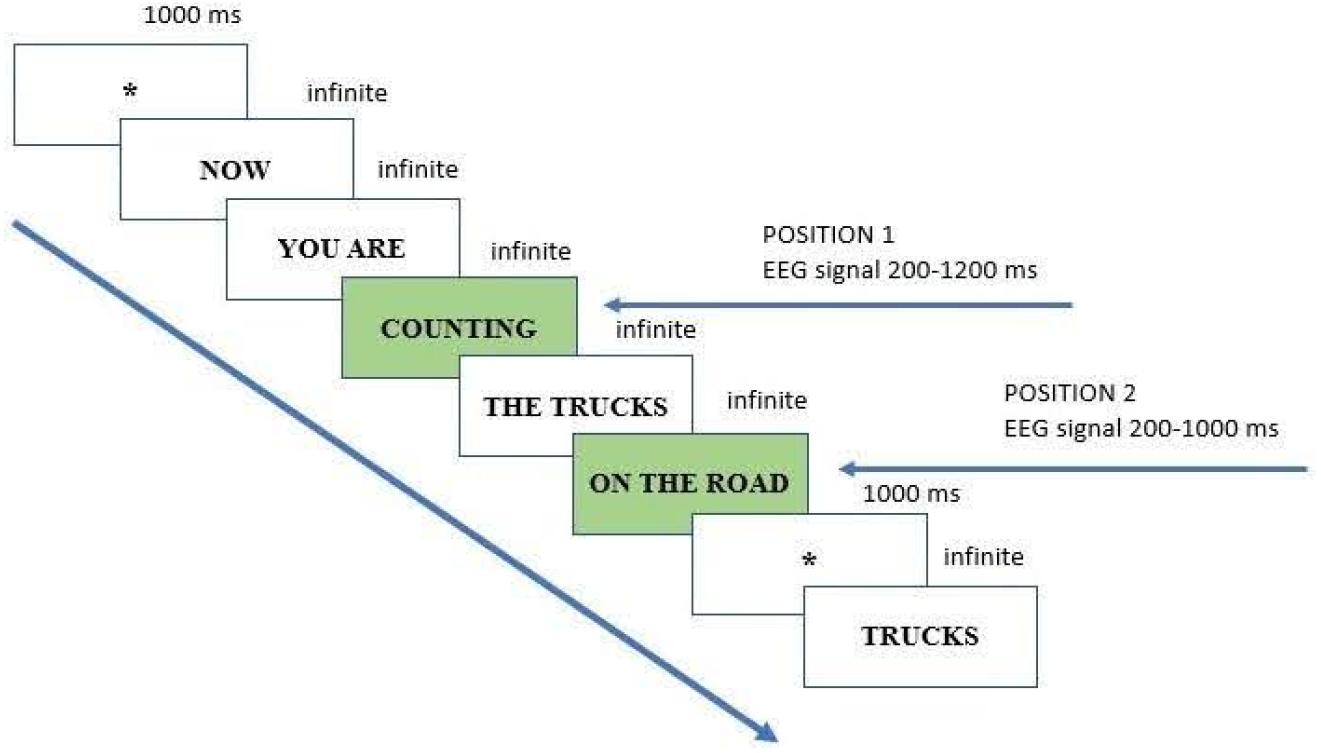
Time sequence of an assay

In each experimental sentence, two loci were used to record the event-related potentials (ERPs) of a word (gerund/participle), corresponding to a central action verb (POSITION 1) and two syntagmas (direct object and adverbial), related to final phrases (POSITION 2).

The purpose of recording both positions was to check the time course of the effort level processing in contexts of different abstraction levels throughout the sentence. To analyze the electrophysiological signal at POSITION 1, 200 ms prior to the analyzed position were designated as the baseline, and the total recording window was extended to 1200 ms after the presentation of the stimulus. For POSITION 2, 200 ms were also designated as the baseline, but the time window was extended to 1000 ms.

The experiment was self-administered using E-Prime 2.0 Professional software [42].

### 2.4. Electrophysiological Record

A high-density EEG setup was used, with 58 thin Ag/AgCl electrodes mounted on an elastic cap that fitted the size of the head. Five electrodes were of the cup-type (10 mm in diameter) and were placed in the ocular area: one under the left eye and another next to the ridge of the right eye to measure eye movements. Two electrodes were placed in the mastoid area (under each ear) and were used as an average reference for the rest of the electrodes (monopolar recording), and one on the forehead was used as grounding.

Electrode locations on the scalp were: FP1, FP2, F3, F4, C3, C4, P3, P4, O1, O2, F7, F8, T7, T8, P7, P8, FZ, CZ, PZ, F1, F2, P1, P2, AF3, AF4, P5, P6, FC5, FC6, C5, C6, TP7, TP8, PO5, PO6, FPZ, FCZ, CPZ, POZ, OZ, PO3, PO4, CP1, CP2, CP3, CP4, C1, C2, F5, F6, FC3, FC4, FC1, FC2, CP5, CP6, PO7, and PO8. These locations followed the standard 10/20 electrode location system. The inter-electrode impedance was kept below 5 kΩ. The biosignals were processed by a Neuronic amplifier in a band between 0.02–100 Hz.

In the ERP analysis, all windows that contained eye movements (EOG greater than 80 *µ*V) and artifacts that contaminated the measurements, such as noise from body movement or facial muscles, among others, were eliminated from the individual averages. This procedure was conducted using the filter provided by the Neuronic analysis program, which performs automatic filtering. However, each window was then manually corrected to ensure that the elimination criteria were consistently applied, allowing for stricter cleaning of each window.

The stimuli for the EEG recording system were presented using E-Prime 2.0 Professional software [42]. The EEG signals were prepared for ERP analysis to determine whether endogenous components, such as the N400, distinguished between factual and counterfactual meanings, as well as between high and low levels of mental effort. The choice of time windows for the ERP/EEG analysis was based on theoretical and statistical criteria. Specifically, the nonparametric statistical method of permutations (included in the Neuronic EP Workstation package) was applied to estimate pairwise t-test comparisons between the levels of a variable at each data point [43]. The ERP segments that met the statistical criteria in the permutation tests were selected as time windows for analysis, according to the topographic distribution of the ERP components described in the literature.

## 3. Results

### 3.1. Position 1

#### 3.1.1. 200-600 ms Window

There were significant effects on the set of 58 electrodes with a significant interaction between Context and Effort, *F* (1, 25) = 4.476, MSe = 4384465334.7, 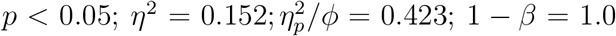 (Greenhouse-Geisser correction). In the first position, the N400 component is expected, characterized by a negativity between 200 and 600 ms, being greater in central-parietal sites with a bias towards the right hemisphere for written language [44]. Older adults exhibit greater N400 priming effects, suggesting enhanced semantic processing, which could explain this lower latency or greater N400 precocity [45].

Thirty-four channels were chosen as the region of interest (ROI): C3, C4, P3, P4, P7, P8, P1, P2, P5, P6, C5, C6, PO5, PO6, PO3, PO4, CP1, CP2, CP3, CP4, C1, C2, CP5, CP6, PO7, PO8, Fz, Cz, Pz, FPz, FCz, CPz, POz, and Oz. A significant interaction was found between the factors Context and Effort, *F* (1, 25) = 4.207, MSe = 330401238.1, 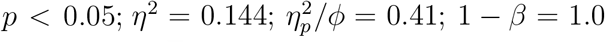 (Greenhouse-Geisser correction). This effect is shown in Figure 2.

**Figure 2.**
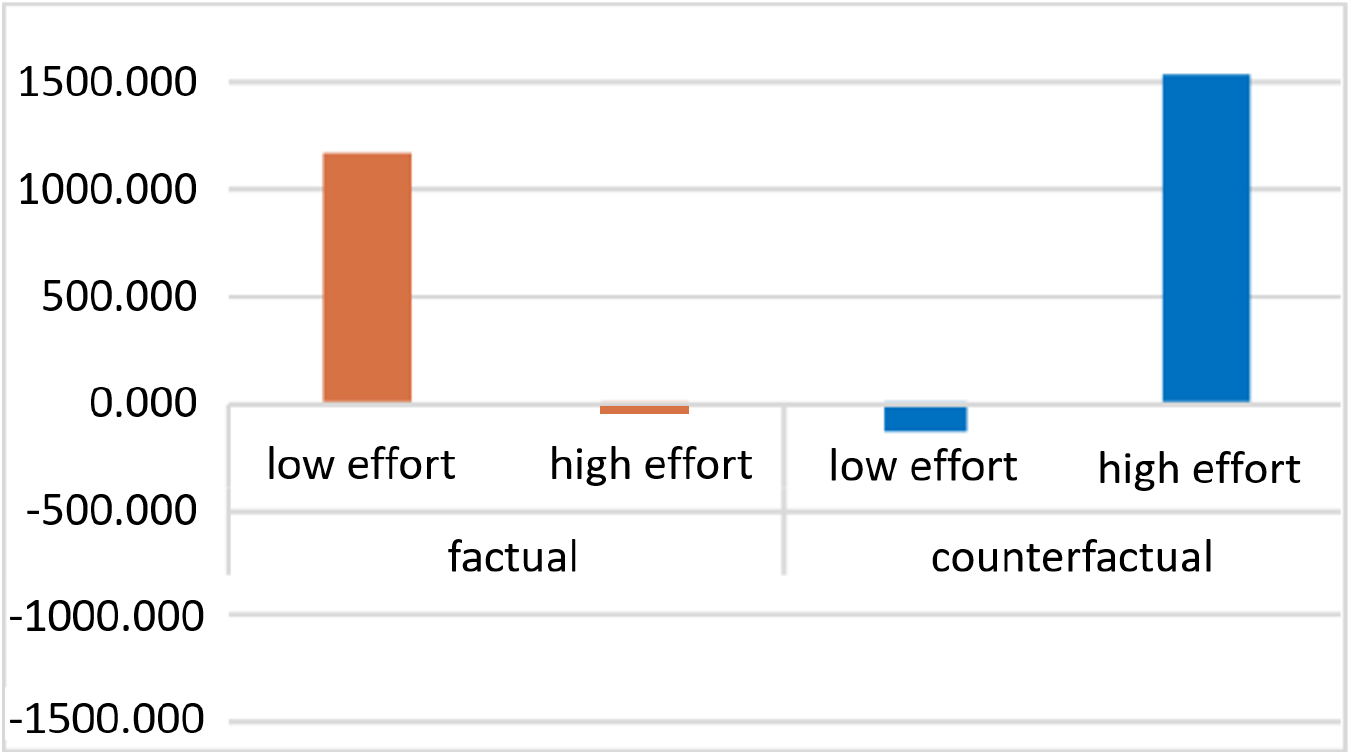
Average ROI 200-600 ms.

The medial channels were analyzed separately, revealing a significant interaction between Context and Effort, *F* (1, 25) = 4.238, MSe = 63211215.87, 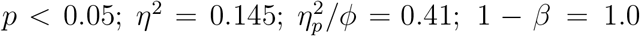 (Greenhouse-Geisser correction). In this time window, the N400 was primarily expressed in the posterior parietal and central areas, and to a lesser extent in the parieto-occipital areas.

Figure 3 shows the voltage differences (amplitude) between the different experimental conditions. The positivity of the high mental effort counterfactual predominates, followed by the low mental effort factual. The topographic distribution of this temporal window is mainly observed in the right central parietal areas according to the topography of a classic N400. However, the brain wave morphology shows several voltage changes within the temporal range, which is consistent with the typical waveforms observed in the study population.

**Figure 3.**
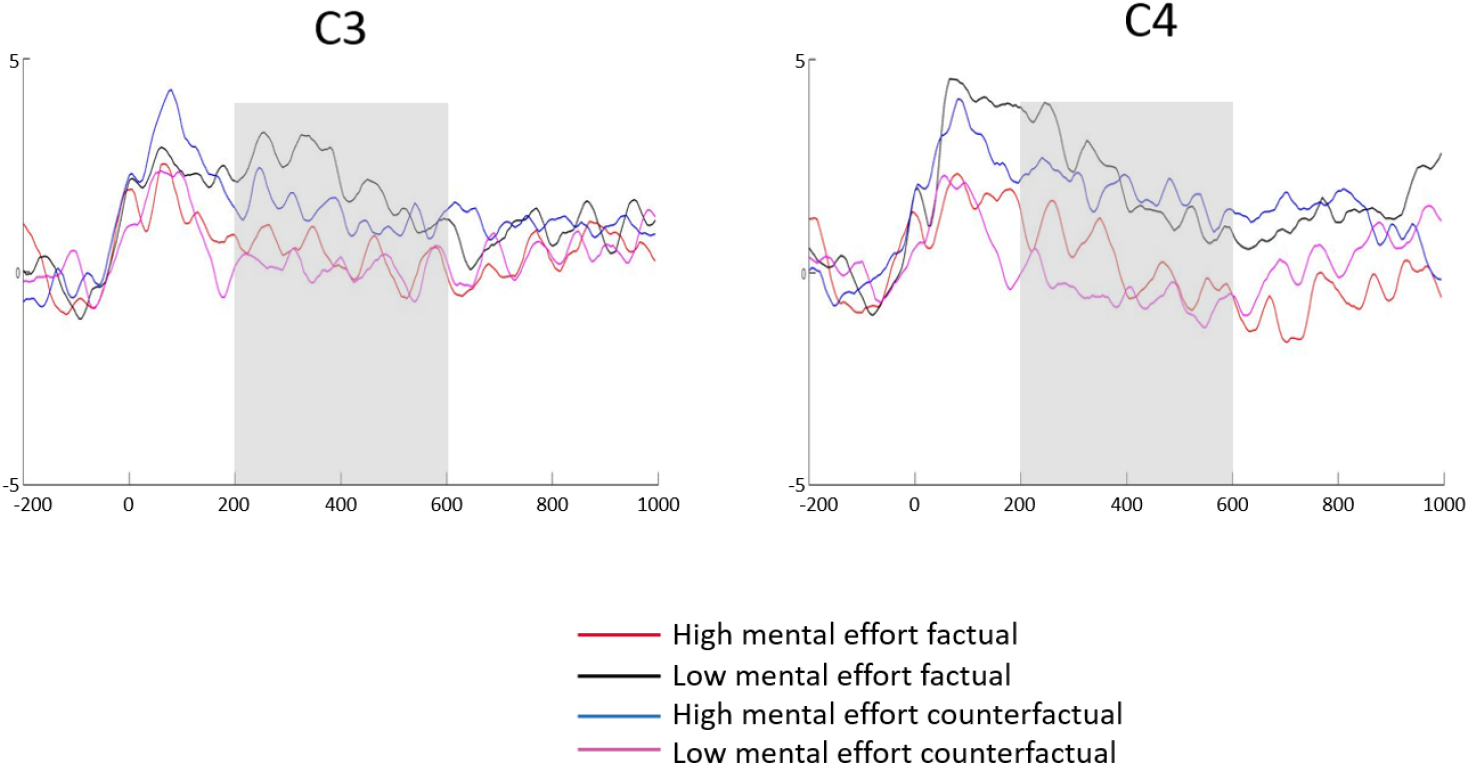
Experimental conditions at C3 and C4 electrodes between 200-600 ms.

#### 3.1.2. 550-700 ms Window

Significant effects were observed on the set of 58 electrodes, with a significant interaction between Context and Effort, *F* (1, 25) = 4.119, MSe = 431735364.2 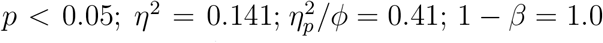 (Greenhouse-Geisser correction). This is depicted in Figure 4.

**Figure 4.**
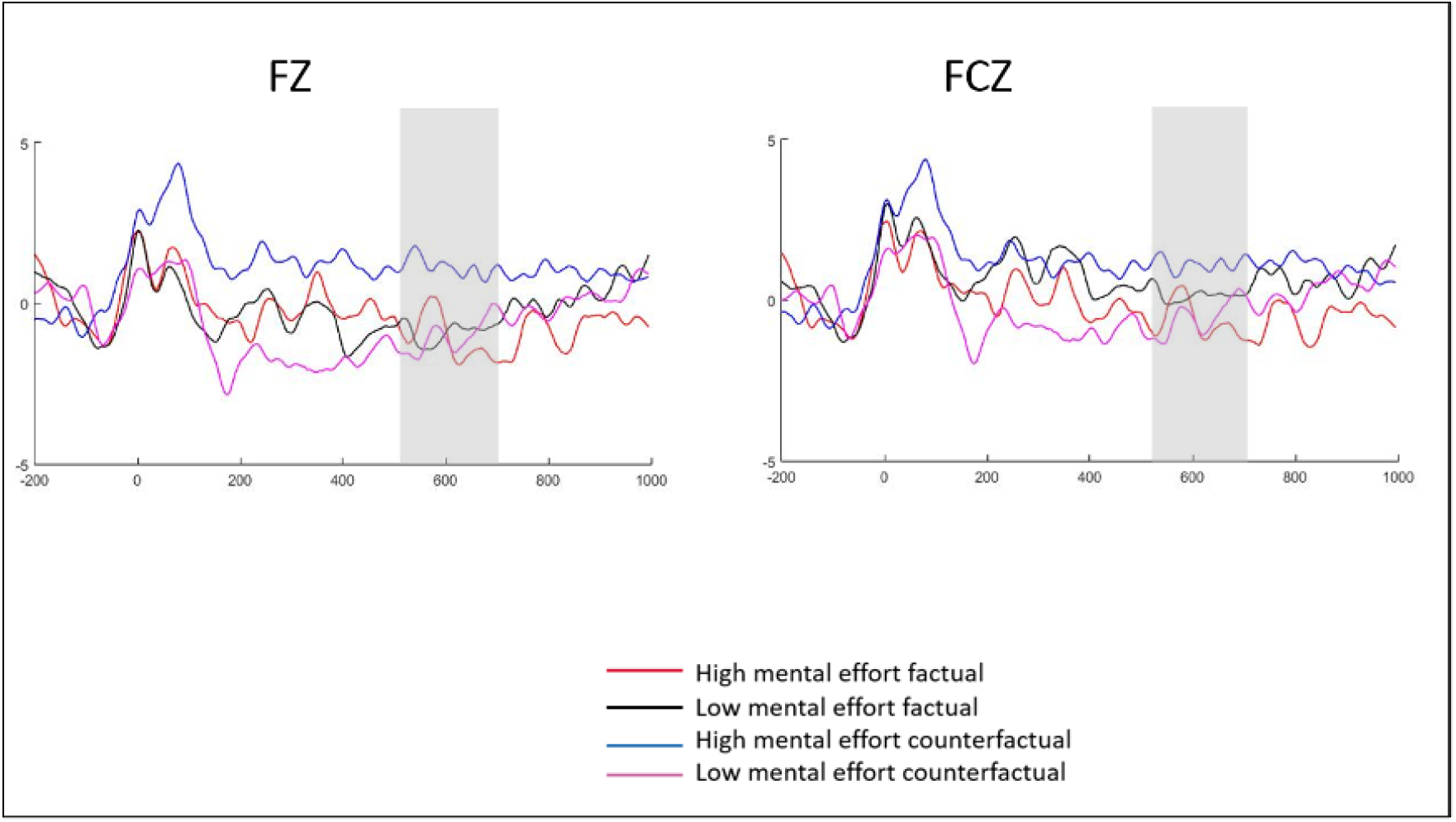
Experimental conditions at the Fz y FCz between 550-700 ms.

Following [46], who studied whether the P600 component reflects only syntactic or also semantic/pragmatic processes, 40 electrodes were analyzed: Fz, FCz, Cz, CPz, Pz, POz, F3, F4, C3, C4, P3, P4, O1, O2, F7, F8, T7, T8, P7, P8, P5, P6, FC5, FC6, C5, C6, TP7, TP8, PO3, PO4, CP3, CP4, F5, F6, FC3, FC4, CP5, CP6, PO7, and PO8. A significant interaction was found between Context and Effort, *F* (1, 25) = 4.261, 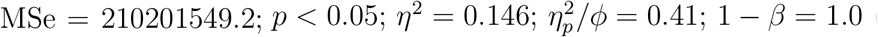 (Greenhouse-Geisser correction). In parallel, a significant interaction was observed in the medial channels: Fz, FCz, Cz, CPz, Pz, and POz, *F* (1, 25) = 4.207, 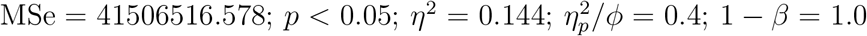 (Greenhouse-Geisser correction).

### 3.2. Position 2

#### 3.2.1. 630-750 ms Window

The medial channels (Fz, Cz, Pz, FPz, FCz, CPz, POz, and Oz) were analyzed separately, yielding a significant interaction between Context and Effort, *F* (1, 25) = 6.293, 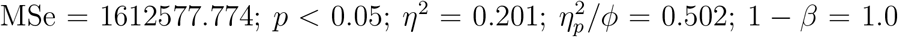 (Greenhouse-Geisser correction). The distribution of the effect in the midline channels indicates greater positivity primarily for central zones and secondarily in frontocentral zones. Similarly, an interesting result was found in the medial channels, showing a significant interaction between Context and Effort, *F* (1, 25) = 6.293, 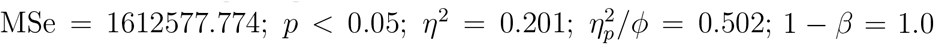 (Greenhouse-Geisser Correction). The data indicate the presence of a focused and limited late positivity that seems to respond mostly to counterfactuals with high mental effort semantics. The results are shown in Figures 5 and 6.

**Figure 5.**
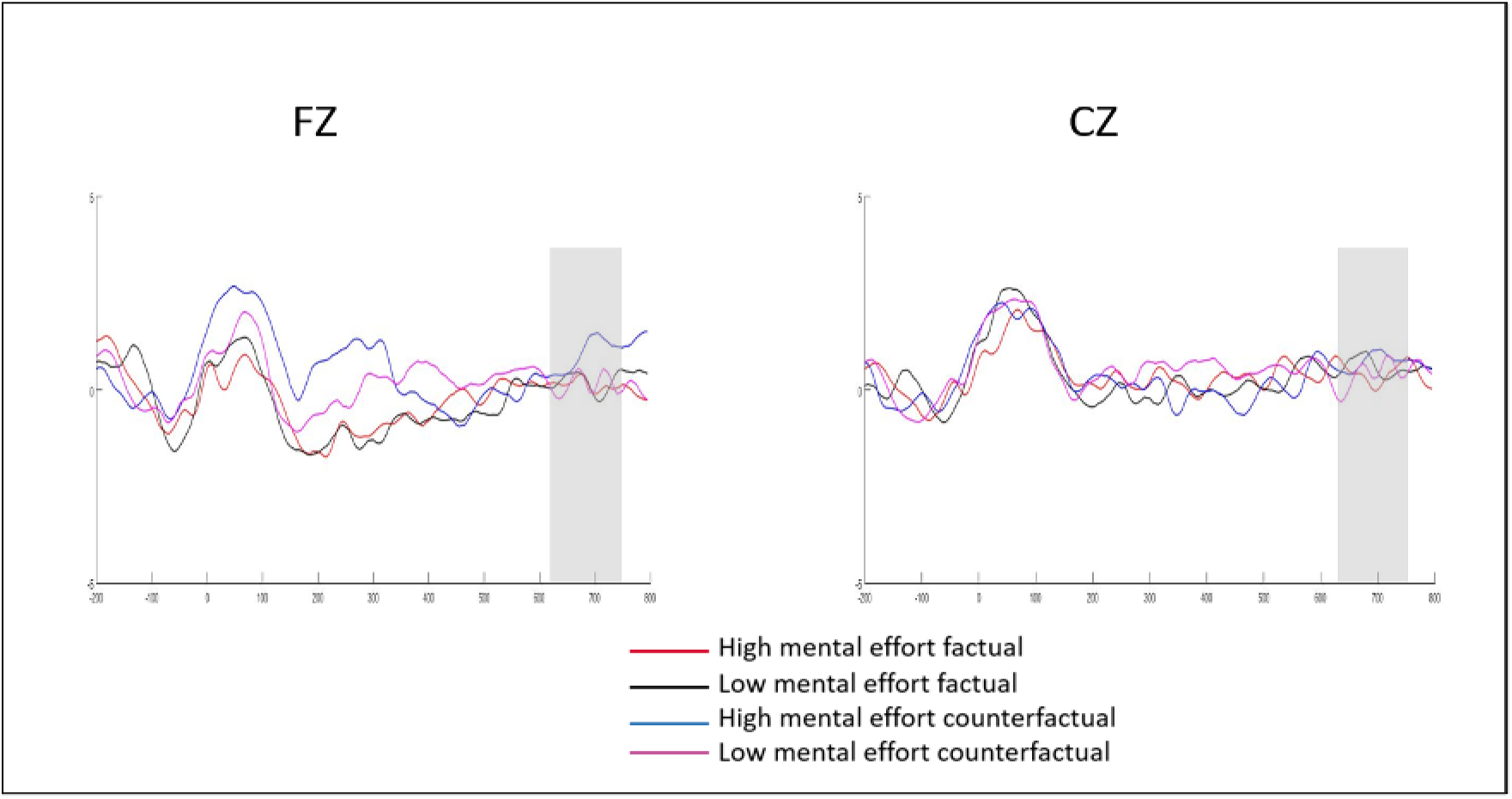
Experimental conditions at the Fz and Cz electrodes between 630-750 ms.

**Figure 6.**
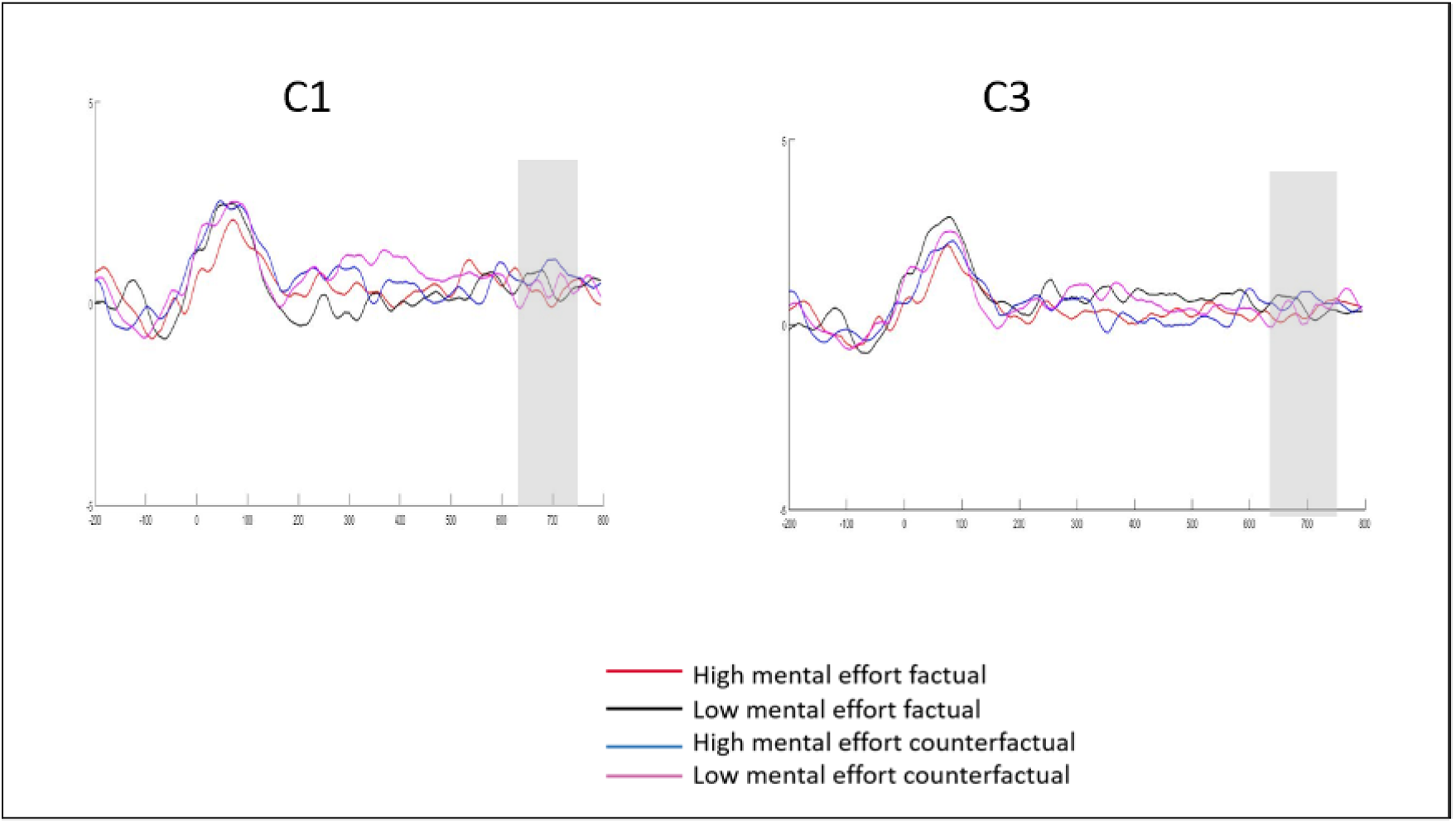
Experimental conditions at the C1 and C3 electrodes between 630-750 ms.

## 4. Discussion

In this study, we explored the ERP components that occur when processing sentences with different levels of mental effort in factual and counterfactual linguistic contexts in the elderly population. The conceptual discussion of these results was carried out under the paradigm of embodied and situated cognition [47, 48].

### 4.1. N400 Component

The elicitation of the N400 ERP component across the three-time windows necessitates an integrated and sequential discussion starting from Position 1. The most striking aspect of this component lies in its early onset and extended latency. The negativity began at 200 ms and continued until 600 ms, revealing an earlier onset by approximately 180 ms and an extension of 160 ms compared to the typical N400 time window observed in younger individuals. This extended latency may reflect the increased size of semantic networks associated with a more extensive lexical-conceptual volume in the elderly [49]. The linguistic features here appear to be shaped by the sensorimotor information linked to the degree of mental effort, with low-effort factual sentences producing greater negativity than high-effort ones. Similarly, low-effort counterfactuals also exhibited greater negativity compared to high-effort counterfactuals.

These findings suggest that the N400 component was modulated early and predominantly by both the level of mental effort and the counterfactual context, integrating corporeal, emotional, and linguistic aspects in early processing stages. The early onset of N400 in older adults could be explained by the increased N400 priming effects observed in this population, indicating enhanced semantic processing [45]. This suggests that older adults may engage in more extensive and earlier semantic processing when faced with varying levels of mental effort and counterfactuality.

One explanation for these results could be that older adults mentally simulate high-effort actions as more challenging, requiring more cognitive resources for proper execution. Conversely, low-effort factual sentences seem to offer greater opportunities for simulation, representation, and semantic integration, making them easier to process. The high-effort counterfactual sentences, however, evoked a more positive response, possibly due to the stimulation of imagining challenging past scenarios, which might be experienced as a sense of challenge accompanied by positive emotions. This positivity may be linked to the emotional salience of these sentences.

Regarding the brain regions involved, the early N400 effect was modulated by the right central and parietal areas, the “standard” N400 by frontal and central areas, and the late N400 by frontal, frontocentral, central, centroparietal, and parieto-occipital regions.

The greater negativity of the high mental effort factual would imply that by progressively building the situation model [50, 51, 52] of the sentence and reaching the critical area where the gerund connects with the direct object, the high effort in indicative grammatical mode tries to recruit personal dispositions that the person unconsciously rejects or does not prefer, which would be interpreted as implausibility or semantic incoherence. The opposite is true for the high mental effort counterfactual, which continues to be salient probably due to emotional and social factors [53, 54]. The social aspect would be explained because the mandate contained in the sentence beginning with “You should have...” is ultimately a social mandate oriented to “should be” that comes from moral standards introjected from the culture and that could eventually imply the review of the conduct from the perspective of other people who judge or evaluate the conduct.

These findings are consistent with studies suggesting that older adults process more concrete elements more efficiently when these elements align with their daily life experiences. The increased salience of high-effort counterfactuals may be attributed to their emotional load and dual semantic representation [55, 56].

### 4.2. Late Positivity

In Position 2, a late positivity was observed, centered on a specific region of interest, reflecting processes of review, re-evaluation, and closure of the global situation model. This late positivity found at the end of sentences in both factual and counterfactual contexts represents an original contribution to the literature as it has not been previously reported. We hypothesize that this positivity is associated with working memory processes, as participants needed to prepare for a subsequent recognition task. This preparation likely required the allocation of more attention and cognitive resources to more abstract elements, particularly those involving high mental effort and counterfactual reasoning.

It is also possible that this working memory is linked to linguistic aspects associated with the double cognitive processing of the counterfactual, which generates a greater cognitive load.

## 5. Conclusion

The present study provides empirical evidence on the temporal dynamics of language processing in the aging brain. Linguistic stimuli, both factual and counterfactual, along with the embodied aspect of mental effort, are found to modulate processing, with each exerting varying degrees of influence over different temporal stages. Counterfactuality appears to facilitate the processing of high-effort language, whereas the interaction between low-effort and counterfactual language produces a negativity associated with the N400 component. A similar dynamic is observed with high-effort factual language, where an early manifestation of the N400 is notable. This early N400 response may be attributed to the elderly’s greater lexical knowledge and compromised inhibitory processes, phenomena apparently stimulated by semantic complexity and mental effort.

Among older adults, increased N400 responses are observed for high-effort factual language compared to their response to low-effort factual language. This observation is in line with evidence suggesting that older people process the corporeal aspects of sentences less easily when they perceive them as implausible or incongruent with their life context. In contrast, counterfactual sentences show a different response pattern in which the initial positivity gradually decreases as events are reinterpreted, thus favoring the factual interpretation. This effect may be due to linguistic-semantic considerations probably influenced by emotional factors, which also embody an essential aspect of language processing.

Our analysis implies that age-related declines in cognitive functions, particularly those associated with the mirror neuron system [57], spanning the inferior frontal lobe, inferior parietal, and superior temporal areas, have implications for several cognitive processes, such as action observation and prediction, theory of mind, empathic resonance, motor resonance, and sensorimotor simulation in older people [58, 59]. The clinical significance lies in that older people, due to decreased activity of this system, may have difficulties understanding and predicting actions described in language, which affects communication and social interaction. This difficulty could lead to smaller social networks, fewer positive social exchanges, less frequent participation in social activities, and higher levels of loneliness, potentially increasing their risk of mental and physical health problems [60, 61], particularly depression. Consequently, it will be essential to develop rehabilitation programs that consider these declines. Interventions that strengthen the activity of the mirror neuron system through action observation exercises and sensorimotor simulation could improve language comprehension, communication, and empathy in older people [62].

In sum, investigating the interaction between neural mechanisms, language processing, and cognitive aging enriches both theoretical understanding and clinical practice, offering valuable insights for interventions targeting older populations. These efforts hold promise for addressing the communication challenges faced by older people and improving their cognitive well-being. Limitations of this study include the lack of a control and/or comparison group, which restricts our ability to attribute the observed effects to the variables studied, leaving room for alternative explanations. Furthermore, the sample was not representative of the general population of older adults, particularly those with lower educational levels, which may affect cognitive performance and functionality of the mirror neuron system.

Future research should include a control group and a more diverse sample with varying educational levels to improve the validity and generalizability of the study. Addressing these gaps will improve the understanding and development of interventions tailored to a broader range of older adults, thereby increasing the effectiveness of potential support strategies.

## Acknowledgements

We appreciate the funding granted by ANID/PIA/Basal Funds for Centers of Excellence FB0003.

## References

[1] L. Barsalou, Grounded cognition, Annual Review of Psychology 59 (2008) 617–645.

[2] M. De Vega, Revisitando la corporeidad del lenguaje narrativo, Revista Signos 54 (107) (2021) 985–1003. doi:10.4067/S0718-09342021000300985.

[3] A. M. Glenberg, M. P. Kaschak, Grounding language in action, Psychonomic Bulletin & Review 9 (3) (2002) 558–565. doi:10.3758/BF03196313.

[4] N. Hinrichs, M. Foradi, T. Yousef, E. Hartmann, S. Triesch, J. Kaßel, J. Pein, Embodied metarepresentations, Frontiers in Neurorobotics 16 (2022) 836799. doi:10.3389/fnbot.2022.836799.

[5] D. A. Allport, E. Funnell, Components of the mental lexicon, Philosophical Transactions of the Royal Society of London. B, Biological Sciences 295 (1077) (1981) 397–410. doi: 10.1098/rstb.1981.0148.

[6] S. Freud, On aphasia: a critical study, International Universities Press, 1891.

[7] J. Locke, An essay concerning human understanding, Kay & Troutman, 1847.

[8] C. Wernicke, Der aphasische Symptomenkomplex, Cohn & Weigert, 1874.

[9] J. Pelkey, Embodiment and language, WIREs Cognitive Science 14 (5) (2023) e1649. doi:10.1002/wcs.1649.

[10] J. L. Salas-Herrera, M. U. Martínez, R. M. Araneda, M. V. De Vos, Comprensión de oraciones de esfuerzo en jóvenes y adultos mayores desde una perspectiva corpórea, Universitas Psychologica 19 (2020) 1–14. doi:10.11144/Javeriana.upsy19.coej.

[11] C. P. Davis, E. Yee, Is time an embodied property of concepts?, PsyArXiv (2023) 1– 27doi:10.31234/osf.io/sqyaj.

[12] F. Pulvermüller, How neurons make meaning: brain mechanisms for embodied and abstract-symbolic semantics, Trends in Cognitive Sciences 17 (9) (2013) 458–470. doi: 10.1016/j.tics.2013.06.004.

[13] L. D. Reggin, L. E. Gómez Franco, O. V. Horchak, D. Labrecque, N. Lana, L. Rio, G. Vigliocco, Consensus paper: Situated and embodied language acquisition, Journal of Cognition 6 (1) (2023) 63. doi:10.5334/joc.308.

[14] A. Körner, M. Castillo, L. Drijvers, M. H. Fischer, F. Günther, M. Marelli, O. Platonova, L. Rinaldi, S. Shaki, J. P. Trujillo, O. Tsaregorodtseva, A. M. Glenberg, Embodied processing at six linguistic granularity levels: A consensus paper, Journal of Cognition 6 (1) (2023) 60. doi:10.5334/joc.231.

[15] E. Venter, Toward an embodied, embedded predictive processing account, Frontiers in Psychology 12 (2021) 543076. doi:10.3389/fpsyg.2021.543076.

[16] M. Kiefer, F. Pulvermüller, Conceptual representations in mind and brain: Theoretical developments, current evidence and future directions, Cortex 48 (7) (2012) 805–825.

[17] C. L. Moody, S. P. Gennari, Effects of implied physical effort in sensory-motor and pre-frontal cortex during language comprehension, NeuroImage 49 (1) (2010) 782–793. doi:10.1016/j.neuroimage.2009.07.065.

[18] M. Urrutia, M. De Vega, Language and action: A current revision to embodiment theories, RLA. Revista de Lingüística Teórica y Aplicada 50 (1) (2012) 39–67. doi: 10.4067/S0718-48832012000100003.

[19] L. Miles, L. Nind, C. Macrae, Moving through time, Psychological Science 21 (2010) 222–223.

[20] R. H. Desai, J. R. Binder, L. L. Conant, Q. R. Mano, M. S. Seidenberg, The neural career of sensory-motor metaphors, Journal of Cognitive Neuroscience 23 (9) (2011) 2376–2386. doi:10.1162/jocn.2010.21596.

[21] S. A. Beauprez, C. Bidet-Ildei, Perceiving a biological human movement facilitates action verb processing, Current Psychology 38 (5) (2019) 1355–1359. doi:10.1007/s12144-017-9694-5.

[22] E. Portegijs, L. Karavirta, M. Saajanaho, T. Rantalainen, T. Rantanen, Assessing physical performance and physical activity in large population-based aging studies: home-based assessments or visits to the research center?, BMC Public Health 19 (1) (2019) 1–16. doi:10.1186/s12889-019-7869-8.

[23] V. Muffato, C. Meneghetti, V. Di Ruocco, R. De Beni, When young and older adults learn a map: The influence of individual visuo-spatial factors, Learning and Individual Differences 53 (2017) 114–121. doi:10.1016/j.lindif.2016.12.002.

[24] M. D. Pham, A. D’Angiulli, M. M. Dehnavi, R. Chhabra, From brain models to robotic embodied cognition: How does biological plausibility inform neuromorphic systems?, Brain Sciences 13 (9) (2023) 1316. doi:10.3390/brainsci13091316.

[25] M. A. Just, S. D. Newman, T. A. Keller, A. McEleney, P. A. Carpenter, Imagery in sentence comprehension: an fmri study, NeuroImage 21 (1) (2004) 112–124. doi: 10.1016/j.neuroimage.2003.08.042.

[26] M. De Vega, M. Urrutia, B. Riffo, Cancelling updating in the comprehension of counterfactuals embedded in narratives, Memory & Cognition 35 (6) (2007) 1410–1421.

[27] F. Paas, J. E. Tuovinen, H. Tabbers, P. W. Van Gerven, Cognitive load measurement as a means to advance cognitive load theory, Educational Psychologist 38 (1) (2003) 63–71.

[28] M. A. Just, P. A. Carpenter, A. Miyake, Neuroindices of cognitive workload: Neuroimaging, pupillometric and event-related potential studies of brain work, Theoretical Issues in Ergonomics Science 4 (1-2) (2003) 56–88.

[29] S. Weber, A. Aleman, K. Hugdahl, Involvement of the default mode network under varying levels of cognitive effort, Scientific Reports 12 (1) (2022) 6303. doi:10.1038/s41598-022-10289-7.

[30] Y. Ke, T. Jiang, S. Liu, Y. Cao, X. Jiao, J. Jiang, D. Ming, Cross-task consistency of electroencephalography-based mental workload indicators: Comparisons between power spectral density and task-irrelevant auditory event-related potentials, Frontiers in Neuroscience 15 (2021) 703139. doi:10.3389/fnins.2021.703139.

[31] G. F. Wilson, C. R. Swain, P. Ullsperger, Erp components elicited in response to warning stimuli: the influence of task difficulty, Biological Psychology 47 (2) (1998) 137–158. doi:10.1016/s0301-0511(97)00021-5.

[32] E. P. Shaw, J. C. Rietschel, B. D. Hendershot, A. L. Pruziner, M. W. Miller, B. D. Hatfield, R. J. Gentili, Measurement of attentional reserve and mental effort for cognitive workload assessment under various task demands during dual-task walking, Biological Psychology 134 (2018) 39–51. doi:10.1016/j.biopsycho.2018.01.009.

[33] L. E. Ismail, W. Karwowski, Applications of eeg indices for the quantification of human cognitive performance: A systematic review and bibliometric analysis, PLoS ONE 15 (12) (2020) e0242857. doi:10.1371/journal.pone.0242857.

[34] R. M. Byrne, The rational imagination: How people create alternatives to reality, MIT Press, 2005.

[35] H. J. Ferguson, Eye movements reveal rapid concurrent access to factual and counterfactual interpretations of the world, Quarterly Journal of Experimental Psychology 65 (5) (2012) 939–961.

[36] E. Kulakova, M. Aichhorn, M. Schurz, M. Kronbichler, J. Perner, Processing counterfactual and hypothetical conditionals: An fmri investigation, NeuroImage 72 (2013) 265–271.

[37] M. S. Nieuwland, A. E. Martin, If the real world were irrelevant, so to speak: The role of propositional truth-value in counterfactual sentence comprehension, Cognition 122 (1) (2012) 102–109. doi:10.1016/j.cognition.2011.09.001.

[38] J. Albayay, P. Villarroel-Gruner, C. Bascour-Sandoval, V. Parma, G. Gálvez-García, Psychometric properties of the spanish version of the edinburgh handedness inventory in a sample of chilean undergraduates, Brain and Cognition 137 (2019) 103618. doi: 10.1016/j.bandc.2019.103618. URL https://doi.org/10.1016/j.bandc.2019.103618

[39] M. F. Folstein, L. N. Robins, J. E. Helzer, The mini-mental state examination, Archives of General Psychiatry 40 (7) (1983) 812. doi:10.1001/archpsyc.1983.01790060110016.

[40] P. Quiroga, C. Albala, G. Klaasen, Validación de un test de tamizaje para el diagnóstico de demencia asociada a edad, en chile, Revista Médica de Chile 132 (4) (2004) 467–478. doi:10.4067/S0034-98872004000400009.

[41] J. A. Yesavage, T. L. Brink, T. L. Rose, O. Lum, V. Huang, M. Adey, V. O. Leirer, Development and validation of a geriatric depression screening scale: A preliminary report, Journal of Psychiatric Research 17 (1) (1982) 37–49. doi:10.1016/0022-3956(82)90033-4.

[42] W. Schneider, A. Eschman, A. Zuccolotto, E-Prime 2.0, Psychology Software Tools, Inc., 2024. URL http://www.pstnet.com

[43] A. Khosla, P. Khandnor, T. Chand, A comparative analysis of signal processing and classification methods for different applications based on eeg signals, Biocybernetics and Biomedical Engineering 40 (2) (2020) 649–690. doi:10.1016/j.bbe.2020.02.002.

[44] M. F. Issa, Z. Juhasz, G. Kozmann, Eeg analysis methods in neurolinguistics: A short review, Évfolyam 2 (2018) 48–54.

[45] H. O. Tiedt, F. Ehlen, F. Klostermann, Age-related dissociation of n400 effect and lexical priming, Scientific Reports 10 (2020) 20291. doi:10.1038/s41598-020-77116-9.

[46] S. Regel, L. Meyer, T. C. Gunter, Distinguishing neurocognitive processes reflected by p600 effects: Evidence from erps and neural oscillations, PLoS ONE 9 (5) (2014) e96840. doi:10.1371/journal.pone.0096840.

[47] T. Schilhab, Derived embodiment in abstract language, Springer, 2017.

[48] B. Winter, Sensory linguistics: Language, perception and metaphor, John Benjamins Publishing Company 20 (2019).

[49] M. Joyal, C. Groleau, C. Bouchard, M. A. Wilson, S. Fecteau, Semantic processing in healthy aging and alzheimer’s disease: A systematic review of the n400 differences, Brain Sciences 10 (11) (2020) 770. doi:10.3390/brainsci10110770.

[50] D. Morrow, J. Chin, A process-knowledge approach to supporting self-care among older adults, Psychology of Learning and Motivation 77 (2022) 165–191.

[51] J. F. Zacks, E. C. Ferstl, Discourse comprehension, Neurobiology of Language (2016) 661–673.

[52] R. A. Zwaan, Situation model: Psychological, International Encyclopedia of the Social & Behavioral Sciences (2001) 14137–14141.

[53] S. D. Houlihan, M. Kleiman-Weiner, L. B. Hewitt, J. B. Tenenbaum, R. Saxe, Emotion prediction as computation over a generative theory of mind, Philosophical Transactions of the Royal Society A 381 (2023) 20220047. doi:10.1098/rsta.2022.0047.

[54] N. Van Hoeck, P. D. Watson, A. K. Barbey, Cognitive neuroscience of human counterfactual reasoning, Frontiers in Human Neuroscience 9 (2015) 420. doi:10.3389/fnhum.2015.00420.

[55] G. Buccino, I. Colagè, F. Silipo, P. D’Ambrosio, The concreteness of abstract language: an ancient issue and a new perspective, Brain Structure and Function 224 (4) (2019) 1385–1401. doi:10.1007/s00429-019-01851-7.

[56] C. Scorolli, F. Binkofski, G. Buccino, R. Nicoletti, L. Riggio, A. M. Borghi, Abstract and concrete sentences, embodiment, and languages, Frontiers in Psychology 2 (2011) 227. doi:10.3389/fpsyg.2011.00227.

[57] E. Farina, F. Borgnis, T. Pozzo, Mirror neurons and their relationship with neurodegenerative disorders, Journal of Neuro Research 98 (2020) 1070–1094. doi: 10.1002/jnr.24579.

[58] T. H. Bak, The neuroscience of action semantics in neurodegenerative brain diseases, Current Opinion in Neurology 26 (6) (2013) 671–677.

[59] D. V. Moretti, Involvement of mirror neuron system in prodromal alzheimer’s disease, BBA Clinical (2016) 46–53.

[60] A. D. Palmer, J. T. Newsom, K. S. Rook, How does difficulty communicating affect the social relationships of older adults? an exploration using data from a national survey, Journal of Communication Disorders 62 (2016) 131–146. doi:10.1016/j.jcomdis.2016.06.002.

[61] A. D. Palmer, P. C. Carder, D. L. White, G. Saunders, H. Woo, D. J. Graville, J. T. Newsom, The impact of communication impairments on the social relationships of older adults: Pathways to psychological well-being, Journal of Speech, Language, and Hearing Research 62 (1) (2019) 1–21. doi:10.1044/2018_JSLHR-S-17-0495.

[62] J. G. Douma, K. M. Volkers, J. P. Vuijk, M. H. Sonneveld, R. H. Goossens, E. J. Scherder, The effects of observation of walking in a living room environment, on physical, cognitive, and quality of life related outcomes in older adults with dementia: a study protocol of a randomized controlled trial, BMC Geriatrics 15 (1) (2015) 26. doi: 10.1186/s12877-015-0024-1.

